# Some like it hot: efficiency of the type III secretion system has multiple thermosensitive behaviors in the *Pseudomonas* syringae complex

**DOI:** 10.1101/2024.05.31.596889

**Authors:** E. Caullireau, D. Danzi, V.M. Tempo, M. Pandolfo, C.E. Morris, E. Vandelle

**Affiliations:** Dipartimento di Biotecnologie, Università degli Studi di Verona, I-37134 Verona, Italia; INRAE, Pathologie Végétale, F-84140, Montfavet, France

**Keywords:** bacterial virulence, T3SS, hypersensitive response, molecular plant-pathogen interactions, environmental factors

## Abstract

The *Pseudomonas syringae* complex is an important group of ubiquitous bacteria containing plant pathogenic strains of which many cause damage and economic losses to a wide range of crops. Efforts to elucidate the determinants of host range have focused on the repertoires of effectors in the Type 3 Secretion System (T3SS). However, recently, we showed that the inability of a *P. syringae* pv. *actinidiae* strain to trigger an hypersensitive response (HR) in *A. thaliana* is due to an inefficient T3SS and not to the absence of a recognized effector. In this context, we compared the efficiency of the T3SS of several *P. syringae* strains belonging to different phylogroups. Through statistical analysis, we assessed the temporal dynamics of the induction of ion leakage, an HR indicator, in *A. thaliana*. We revealed that *Pto*DC3000 *avrB* and *Pma*M6 *avrB* consistently triggered a strong HR while other strains induced it at significantly different intensities depending on temperature. Among thermosensitive strains, both low and warm temperature-dependencies for T3SS efficiency were observed, irrespective of the *in-vitro* growth optimum of the strains. Surprisingly, differences were also observed among quasi-clonal strains. These results reveal a strong, strain-specific regulatory role of temperature in effector injection and reinforce the importance of environmental factors in the outcome of plant-bacteria interactions. Moreover, this work highlights the need to study bacterial virulence for a broader set of strains beyond model strains, such as *Pto*DC3000, that are not representative of the whole *P. syringae* complex.

## Introduction

*Pseudomonas syringae* is a complex of gram-negative bacteria that poses a threat to global agriculture with new outbreaks that emerge regularly (Lamichhane et al., 2014; Lamichhane et al., 2015). *P. syringae* has been divided into at least 13 phylogenetic groups (phylogroups, PG), further divided into 23 clades (Berge et al., 2014). During the infection, *P. syringae* relies on an arsenal of virulence factors, though its pathogenesis is predominantly mediated by the type III secretion system (T3SS), a highly conserved needle-like specialized protein secretion machinery, found in numerous gram-negative bacterial pathogens affecting plants and animals. T3SS forms a continuous channel, guiding the translocation of diverse proteinaceous effectors (T3Es) directly into the host cell or apoplast (Büttner and He, 2009; Jin and He, 2001; Li, 2002; Roine et al., 1997). *P. syringae* strains with the canonical T3SS commonly harbor a repertoire of approximately 15-30 T3SEs, which facilitate the invasion of plant host cells, suppress plant immunity, or interfere with essential plant cellular processes (Chang et al., 2005; Cunnac et al., 2009; Galán and Collmer, 1999; Guo et al., 2009; Guttman et al., 2002; Hueck, 1998; Lee, 1997; Lindeberg et al., 2012; Martel et al., 2022; Mudgett and Staskawicz, 1998; Xin et al., 2016). Of note, there is no direct relationship between the size or content of the T3SE repertoire and host range (Baltrus et al., 2017; Morris et al., 2019).

According to the ‘disease triangle’ paradigm (Stevens, 1960), plant diseases arise from the interplay between a susceptible plant and a virulent pathogen, under conducive environmental conditions, resulting in a compatible reaction. Overall plant-pathogen interaction mechanisms are highly influenced by environmental conditions, which can either favor, have neutral effects, or hinder both partners (Velásquez et al., 2018; Xin et al., 2018). Among the environmental factors, temperature has a regulatory role for virulence in plant-associated bacteria (reviewed in references Smirnova et al., 2001; Velásquez et al., 2018).

Within the *P. syringae* species complex, temperature variations impact various traits crucial for bacterial fitness, including phytotoxin expression, exopolysaccharide biosynthesis, motility or the quorum-sensing system (Aguilera et al., 2007; Aguilera et al., 2017; Arvizu-Gómez et al., 2013; Bender et al., 1999; Braun et al., 2009; Hockett et al., 2013; Krishna et al., 2022; Li et al., 2006; Nüske and Fritsche, 1989; Palmer and Bender, 1993; Peñaloza-Vázquez et al., 1997; Scalschi et al., 2022; Smirnova et al., 2001; Ullrich et al., 2000; Weingart et al., 2004). T3SS has also been investigated concerning its possible thermoregulation. In *P. syringae* pvs. *syringae* and *tomato*, the secretion of HopPsyA and AvrPto effectors *in vitro* was impaired at temperatures above 22°C and 20°C, indicating some kind of low-temperature-dependency for their T3SS efficiency (Van Dijk et al., 1999). In contrast, recent findings in *Pto* DC3000 revealed that the translocation of T3Es in *A. thaliana* cells is enhanced at 30°C compared to 23°C, indicating the possible T3SS efficiency even at elevated temperatures (Huot et al., 2017). However, in tomato, the expression of T3SS-related genes *hrpL* and *hrpA* in this same strain was not affected by a temperature rise from 26°C to 31°C, although the expression of the avirulence gene *avrPtoB* was significantly reduced (Scalschi et al., 2022). On the other hand, for *P. syringae* pv. *actinidiae* strains, it was shown that there was no difference in *hrpA1* promoter activity *in vitro* across a temperature range of 18-28°C, nonetheless, substantial differences were evident at a functional level, demonstrating a low-temperature-dependency for T3SS efficiency for these strains (Puttilli et al., 2022). However, the complete implications of warm temperatures on *P. syringae* virulence in plants remain poorly understood (Hockett et al., 2013; Li et al., 2006). Studies indicate complex interactions with plant immune signaling pathways, emphasizing the need for further research to understand their implications in terms of disease (Cheng et al., 2013; Huot et al., 2017).

Evidence of the influence of temperature in regulating virulence factors in *P. syringae* species complex has been documented, however, there have not been studies dedicated to comparing strains under identical conditions. Consequently, we wanted to further understand the role of the temperature on the function/activation dynamics of the T3SS by investigating a broader range of diversity within the *P. syringae* species complex under comparable conditions. For this purpose, we transformed 13 strains of *P. syringae* in phylogroups 1, 2 and 3 to carry the effector AvrB and infiltrated them into *Arabidopsis thaliana* Col-0 leaf disks. In this model plant, AvrB is indirectly recognized by the resistance protein RPM1, through RIN4 modification, which triggers HR (Innes et al., 1993; Mackey et al., 2002; Mudgett, 2005). Infiltrated leaf disks were incubated at different temperatures and the quantification of electrolyte leakage served as the indicator of HR onset. Conductivity increase was used as a proxy for T3SS efficiency, and this allowed us (i) to effectively discriminate strains based on a robust phenotype directly linked to the activation of the T3SS, and (ii) to observe different behaviors in response to temperature.

## Results

### The intensity of HR induction is not strictly reproducible for individual strains

To evaluate T3SS functionality at different temperatures, *avrB*-expressing *P. syringae* strains were infiltrated into *A. thaliana* Col-0 leaf disks. As a T3SS-dependent mechanism, HR was quantified by measuring the increase in ion leakage due to dying cells. Conductivity levels measured among the diverse replicated experiments were highly variable. The conductivity levels measured for a given modality (*i.e.*, one strain infiltrated and incubated at one temperature) doubled or tripled among replicated experiments. Four examples of this variability are shown in Figure 1. For instance, across the seven replicated experiments for strain USA007 *avrB* at 18°C, the ratio of areas under the curves calculated between the one that induced the most and the least was equal to 2.8 (Figure 1b). Moreover, conductivity showed considerable heteroscedasticity, *i.e.* there was a positive correlation between the mean area under the curves and their variance with Spearman’s coefficient = 0.91; p < 0.05 (Figure S1). This large variability raised the question of how to quantify HR induction to compare the strains, as addressed in the next section.

**Figure 1.**
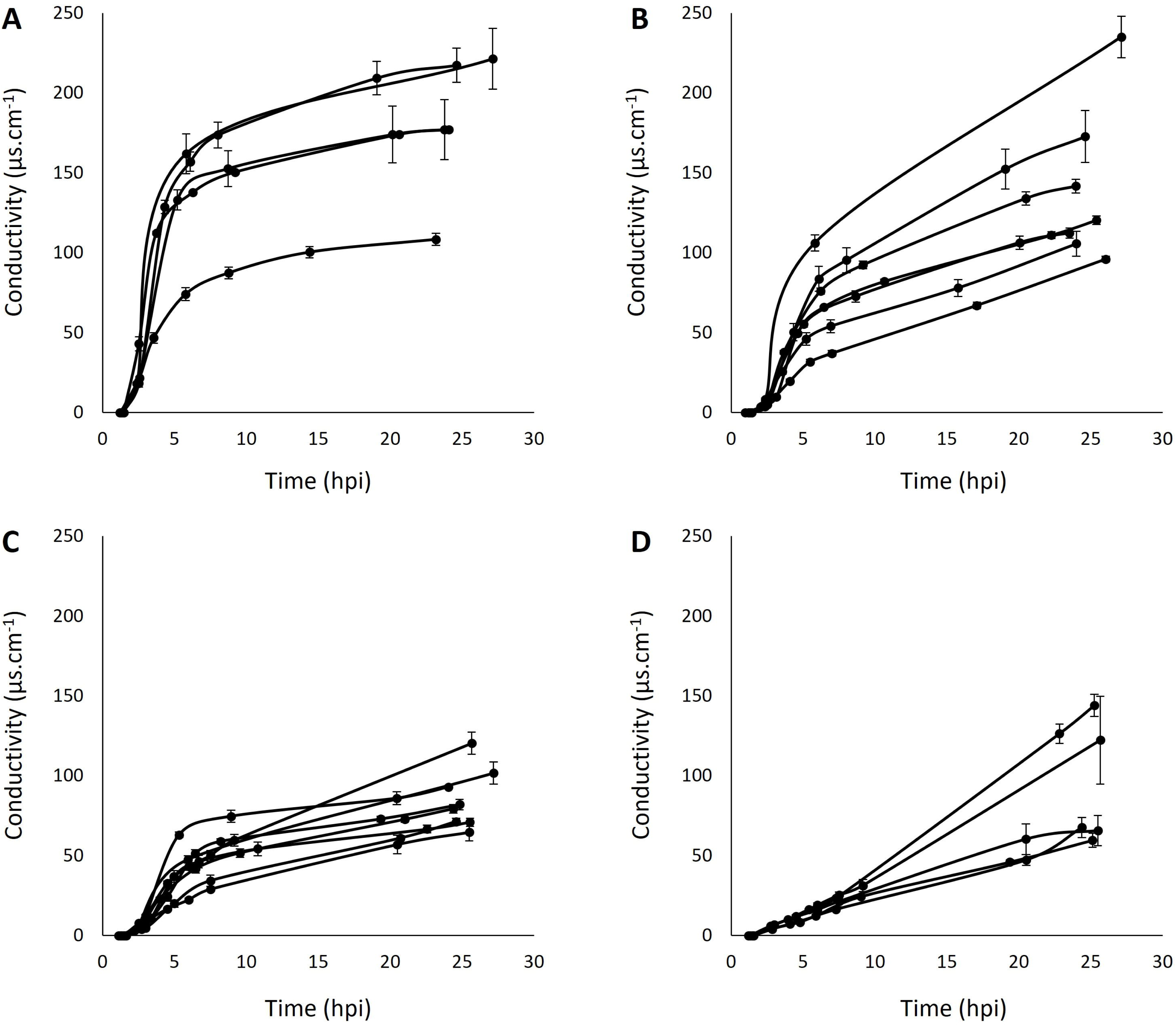
Electrolyte leakage curves obtained post-infiltration for strain M6 *avrB* at 18°C (a), USA007 *avrB* at 18°C (b), LAB0041 *avrB* at 24°C (c) and CC1498 *avrB* at 28°C (d). Each curve represents the mean increase in conductivity measured over time for one independent biological replicate performed with three technical replicates each. Error bars represent standard error. For each curve, the area under the conductivity progress curve (AUCPC) was also calculated, as well as the ratio between the highest and the smallest AUCPC for each strain-temperature couple.

### The intensity of HR induction varies among strains

To compare HR induction intensity among strains, we accounted for the variability described above by fitting Linear Mixed effects Models to our data. This statistical approach allowed us to classify strains as inducers or non-inducers at each of the temperatures that we tested. This revealed that almost all the *P*. *syringae* strains induced HR except for J35 *avrB* at all the temperatures, MAFF302273 *avrB* at 24°C and 28°C, B728a *avrB* at 24°C, and 1448A *avrB* and CRA-FRU 8.43 *avrB* at 28°C (Table 1). The inducer strains led to conductivity levels that were significantly greater than those measured in the mock treatment (10mM MgCl_2_ or *hrp*-inducing medium, HIM) (p < 0.05) according to Linear Mixed-effects Models. More specifically, our results demonstrated that all the inducer strains varied in the intensity of the HR induction they caused. The model estimated the mean difference in points of conductivity measured between the tested modality and the mock treatment. The values presented in Table 1 (“Estimate”) are proxies of the intensity of HR induction for each strain. DC3000 *avrB* and M6 *avrB* displayed the highest values (“Estimate” > 65 regardless of the temperature tested), whereas some others such as the three quasi-clonal strains CC0073 *avrB*, CC0094 *avrB* and CC1498 *avrB* averaged less than 20. These differences are also reflected in the mean areas under conductivity progress curves (AUCPC) at each temperature as shown in Figure 2. This proxy of the HR induction intensity was correlated neither with the *avrB* expression levels (Figure S2) nor with the bacterial growth capacity at a given temperature (Figure S3).

**Figure 2.**
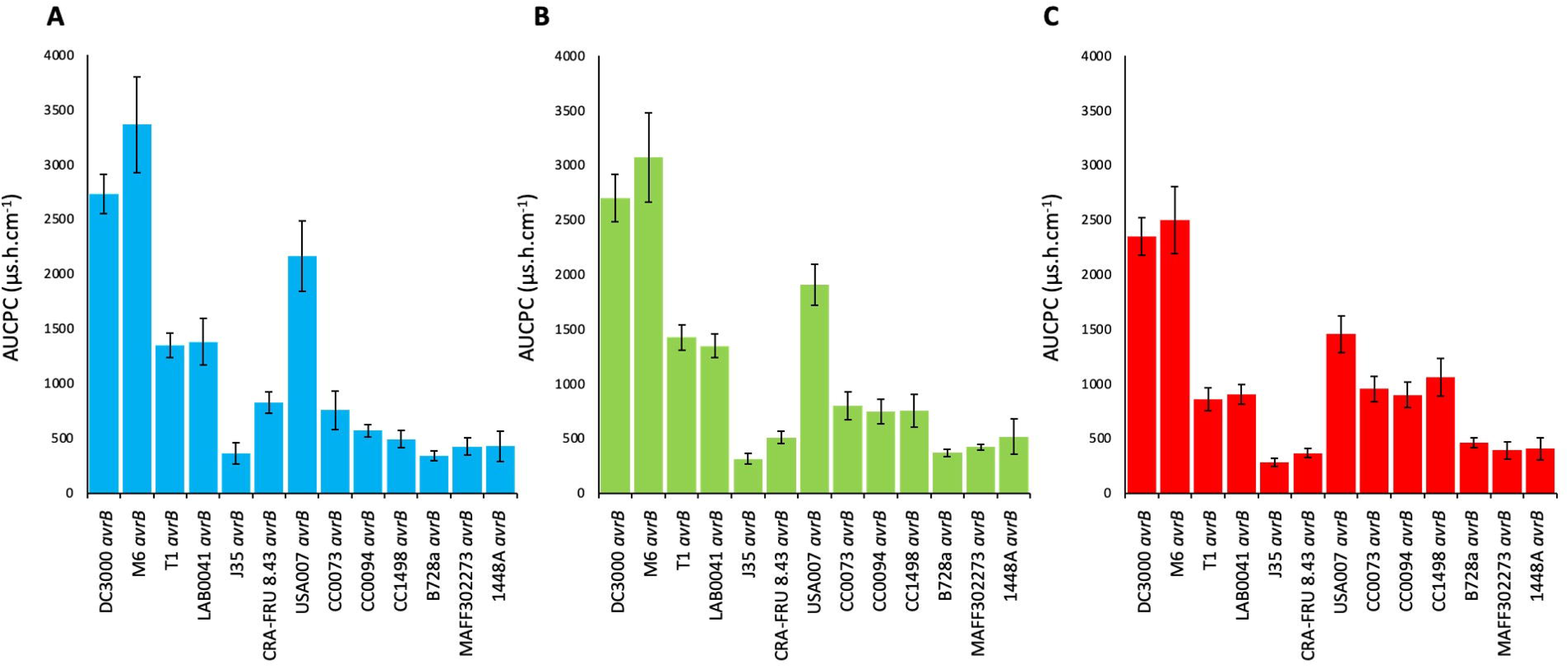
Mean area under conductivity progress curve (AUCPC) calculations for all the replicated experiments (3 ≤ n ≤ 27) of each *avrB*-expressing strain at 18°C (a), 24°C (b) and 28°C (c). Error bars represent standard error.

**Table 1.** Statistics associated with linear mixed-effects models fitted to ion leakage experiment results.

| PHYLOGROUP | STRAIN | INDUCER AT A GIVEN TEMPERATURE |  |  |  |  |  | TEMPERATURE |  |
| --- | --- | --- | --- | --- | --- | --- | --- | --- | --- |
|  |  | 18°C |  | 24°C |  | 28°C |  | *p*-value | *Estimate* |
|  |  | *Estimate* | *p*-value | *Estimate* | *p*-value | *Estimate* | *p*-value |  |  |
| 1a | DC3000 | 69.97 | **2.00E-16** | 73.12 | **1.25E-15** | 65.27 | **8.46E-15** | **8.02E-01** | **-0.19** |
|  | M6 | 95.14 | **2.00E-16** | 89.06 | **5.35E-16** | 73.05 | **1.59E-14** | **1.95E-01** | **-2.07** |
|  | T1 | 28.98 | **2.00E-16** | 29.07 | **2.00E-16** | 13.45 | **2.81E-08** | **1.42E-02** | **-1.26** |
|  | LAB0041 | 31.36 | **4.24E-11** | 29.56 | **2.00E-16** | 16.83 | **2.83E-09** | **5.17E-02** | **-1.22** |
| 1b | J35 | 3.35 | 5.43E-02 | 0.18 | 9.19E-01 | -1.27 | 4.46E-01 | NA |  |
|  | CRA-FRU 8.43 | 14.09 | **4.69E-11** | 4.97 | **4.33E-04** | 0.97 | 4.45E-01 | **3.80E-03** | **-1.28** |
|  | USA007 | 52.24 | **2.00E-16** | 46.69 | **8.58E-16** | 34.15 | **2.26E-12** | **3.35E-02** | **-1.70** |
| 2d | CC0073 | 15.22 | **5.97E-07** | 15.28 | **7.62E-07** | 19.77 | **3.42E-09** | **3.55E-01** | **0.63** |
|  | CC0094 | 6.86 | **6.44E-08** | 9.72 | **2.61E-06** | 14.49 | **8.91E-08** | **9.76E-04** | **0.97** |
|  | CC1498 | 7.89 | **4.96E-06** | 12.69 | **2.24E-05** | 19.96 | **1.52E-08** | **2.58E-02** | **1.36** |
|  | B728a | 3.12 | **2.08E-02** | 2.22 | 1.61E-01 | 4.75 | **8.07E-03** | **4.27E-02** | **0.37** |
|  | MAFF302273 | 3.17 | **1.73E-02** | 2.88 | 9.68E-02 | 1.78 | 3.77E-01 | NA |  |
| 3a | 1448A | 5.14 | **1.07E-02** | 6.73 | **9.73E-03** | 2.46 | 2.41E-01 | **6.41E-01** | **0.53** |

The effect of strains was calculated for each mutant bacterium at each temperature by comparing with the results obtained for the mock treatment (10mM MgCl<sub>2</sub>). Significant values of the estimate parameter ( $p < 0.05$ , in bold) are associated with HR inducing conditions. The temperature effect was evaluated for each *avrB*-expressing strain, inducing HR at least at two temperatures (non-inducer strains are highlighted in grey), comparing the electrolyte leakage curves obtained at these different temperatures. Strains insensitive to the temperature ( $p > 0.05$ ) are highlighted in green, strains leading to minor changes in conductivity levels as the temperature increases (negative estimate, $p < 0.05$ ) are highlighted in blue, and strains displaying a warm-dependency (positive estimate, $p < 0.05$ ) are highlighted in red. The number of replicated experiments for each modality (strain x temperature condition) varied from 3 up to 27.

### The intensity of HR induction is temperature-sensitive in half of the inducer strains

With Linear Mixed-effects Models (LMM), we categorized the HR inducer strains according to the temperature-sensitivity of HR induction. Six out of eleven HR inducer strains were insensitive to a change in temperature (DC3000 *avrB*, M6 *avrB*, LAB0041 *avrB*, CC0073 *avrB*, 1448A *avrB*) (p > 0.05) (Table 1). The other HR inducer strains were temperature-sensitive based on LMM analyses (p < 0.05) and could be categorized into two groups according to whether the “Estimate” values were negative or positive. In one group, strains showed higher conductivity with lower temperatures (T1 *avrB*, CRA-FRU 8.43 *avrB* and USA007 *avrB*). In the case of CRA-FRU 8.43 *avrB* strain, the decrease in conductivity observed with increasing temperature led the strain to be categorized as a non-inducer at 28°C (Table 1). In the other group, strains showed higher conductivity with higher temperatures (CC0094 *avrB*, CC1498 *avrB* and B728a *avrB*). Temperature-sensitivity of the *avrB*-expressing strains was further confirmed by the analysis of HrpZ secretion *in vitro* as an assessment of T3SS functionality (Figure 3).

**Figure 3.**
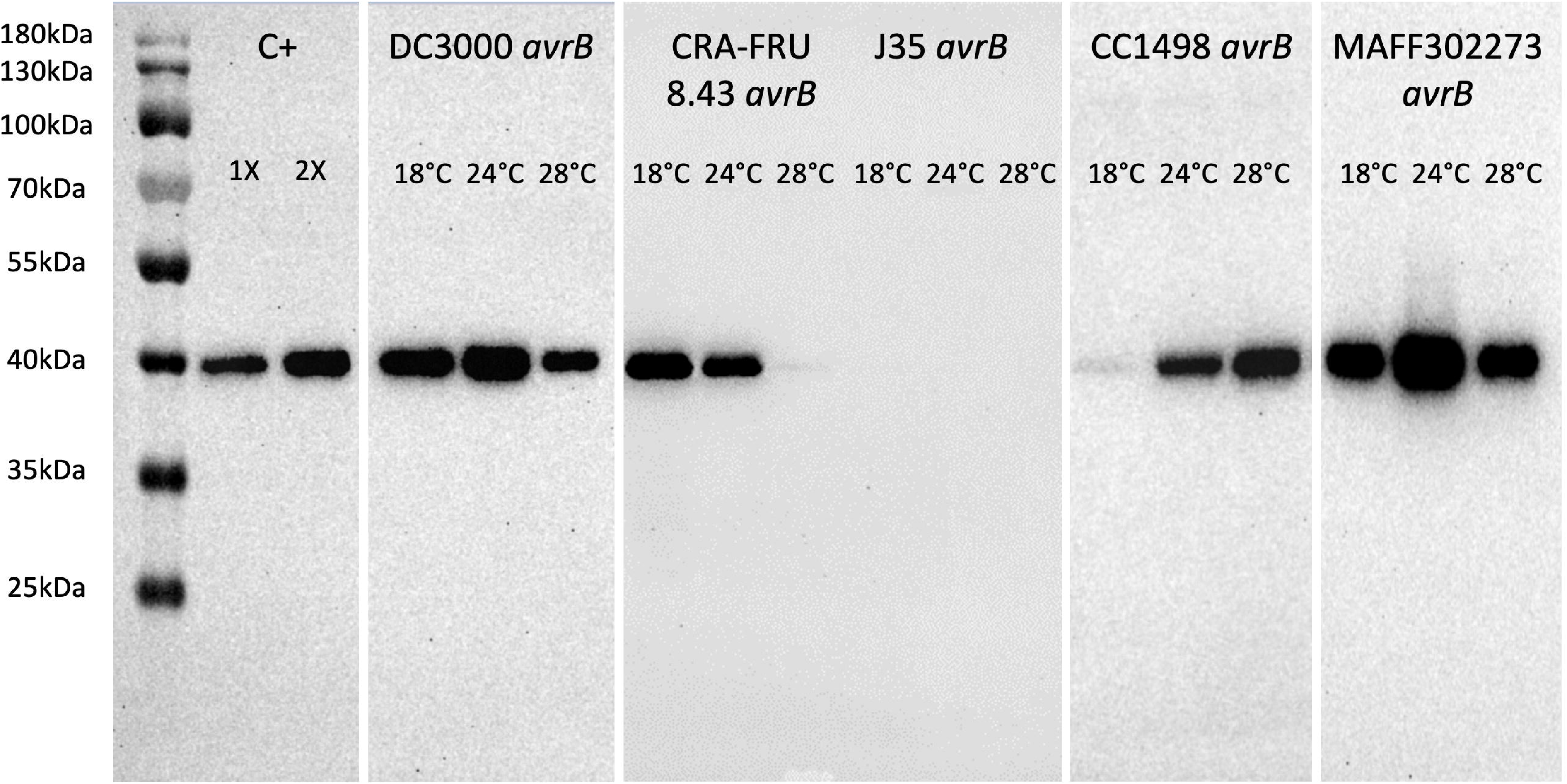
HrpZ secretion profiles across a range of temperature. Strains expressing *avrB* were incubated in *hrp*-inducing medium (HIM) at 18°C, 24°C or 28°C for 5 hours. Cell-free culture supernatant was obtained by centrifugation and filtration. Proteins of bacterial culture supernatant were then precipitated with trichloroacetic acid (TCA) and analyzed by Western blot using an anti-HrpZ antibody. The expected molecular weight is *ca.* 40kDa. A mix of samples proven to contain HrpZ in previous replicates was used as the positive control (C+) at two different concentrations (1X and 2X). Samples were run in different gels that were transferred onto membranes whose pictures were cropped and merged into a single figure.

### The natural genomic background of the WT strains does not change the temperature responsiveness patterns observed with *avrB*-expressing strains

Keeping in mind that the elicitation of the HR by AvrB could be subjected to a variety of factors, including the action of other effectors, both in terms of host responses and competition for delivery by the T3SS, we verified the functionality of the T3SS in vitro, assessing the secretion of the HrpZ protein. While HrpZ detection for J35 *avrB* was consistent with its categorization as an HR non-inducer, strain MAFF302273 *avrB* displayed a functional T3SS at all tested temperatures (Figure 3). Since this strain constantly did not lead to conductivity increase once infiltrated into leaf disks, this may suggest that MAFF302273 contains in its repertoire some suppressive effector(s) blocking the ability of AvrB to trigger the HR in *A. thaliana*. Consequently, we established the effector repertoire composition of all the strains by collecting information already available in the literature for 10 strains and by performing effector prediction in silico for 3 others, following the procedure reported previously (Dillon et al., 2019; Laflamme et al., 2020). However, to the best of our knowledge, no evidence of direct or indirect antagonism between AvrB and other effectors naturally present was found in any of the strains (Table S2). Furthermore, we accounted for the outcome resulting from the interaction between *A. thaliana* Col-0 and our subset of strains in their WT version. Firstly, many of them do not result in HR induction, due to compatible interaction (DC3000, M6) or non-host incompatible interaction without HR (CRA-FRU 8.43, B728a, 1448A), hence justifying the need to introduce *avrB* to study T3SS efficiency with this proxy (Table S3). On the other hand, the presence of *avrB* does not necessarily modify the behavior of the strain. For instance, CC0094 WT induced HR in *A. thaliana* Col-0, displaying the same warm-temperature dependency as observed with the *avrB*-expressing strain (Figure S4a). Likewise, the low-temperature dependency behavior of CRA-FRU 8.43 was conserved across both strain genotypes (with or without *avrB*) in the resistant *Actinidia arguta* (non-host plant inducing effector-mediated HR with strains belonging to the same biovar 3; Figure S4b; Table S3). Altogether, these findings demonstrate that although the HR induction by AvrB in our experiment does not take place in isolation from the background of both interaction partners, the various outcome scenarios with the WT strains allow us to support our conclusions made with the *avrB*-expressing strains, hence confirming the robustness of our approach.

### Temperature-responsiveness patterns vary among strains with trends that are consistent within a given phylogroup

A given phylogroup could contain strains that induced HR in temperature-sensitive and temperature-insensitive ways (Table 1). However, the trend in temperature sensitivity was constant withing a phylogroup among its temperature-sensitive strains. Indeed, the group of strains whose T3SS was more efficient at lower temperatures and the group whose T3SS was more efficient at higher temperatures belong to the phylogroups 1 and 2d, respectively (Table 1). Nevertheless, dramatic differences among strains even those that are very closely related phylogenetically, were still observed as illustrated by the three quasi-clonal *avrB*-expressing strains from phylogroup 2d, CC0073, CC0094 and CC1498 (Table 1) (Figure 4). CC0073 *avrB* was temperature-independent in its ability to induce HR, while CC0094 *avrB* and CC1498 *avrB* both showed clearly greater capacity at warmer temperatures. Moreover, in terms of intensity, the inoculation with CC1498 *avrB* led to higher values of conductivity measured at 24 hpi than CC0073 *avrB* and CC0094 *avrB*. Finally, in terms of kinetics, CC0073 *avrB* was responsible for a faster HR induction at the beginning of the interaction that stalled – no matter the temperature–, whereas CC0094 *avrB* and CC1498 *avrB* were much slower in inducing HR (Figure 4).

**Figure 4.**
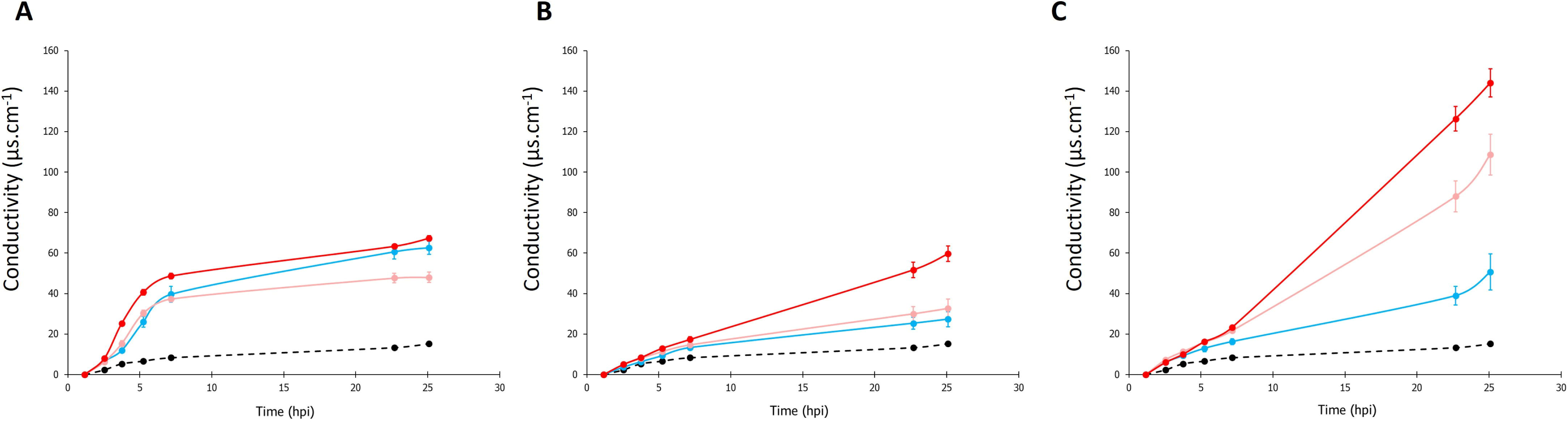
Electrolyte leakage curves over time obtained post-infiltration for CC0073 *avrB* (a), CC0094 *avrB* (b) and CC1498 *avrB* (c). Infiltrated leaf disks were incubated at 18°C (blue), 24°C (pink) and 28°C (red). Conductivity following infiltration of the mock treatment (10mM MgCl_2_) and incubation at 24°C is represented by the black dotted line. Data represent one single representative biological replicate with all strains and temperature conditions evaluated simultaneously with three technical replicates each. Error bars represent standard error.

### Strains from the *P. syringae* complex display diverse speeds of HR induction

To account for dynamic aspects in the behaviors of *avrB*-expressing *P*. *syringae* strains, the time point when half of the total ions (*i.e.* final conductivity value measured at 24 hpi) has leaked was calculated for each replicated experiment for all modalities that were found to be HR-inducing. We used this parameter – hpi_50%_ – as the indicator of HR induction velocity. An analysis of variance of these ratios revealed significant differences among *avrB*-expressing strains (ANOVA1, p < 0.05). DC3000 *avrB* and M6 *avrB* proved to be the fastest HR inducers with half of their total induction already observable before 5 hpi (Figure 5a). Yet again, the diversity of behaviors within phylogroups was revealed, even among the three *avrB*-expressing quasi-clonal strains (Figure 5, post-hoc Dunn’s test showed that CC0073 *avrB* and CC0094 *avrB* and CC1498 *avrB* groups differed significantly at p < 0.05). Furthermore, hpi_50%_ values were compared across the different temperatures overall revealing that higher temperatures correlated with faster electrolyte leakage, consistent with the results observed for bacterial *in-vitro* growth (Figure 5b).

**Figure 5.**
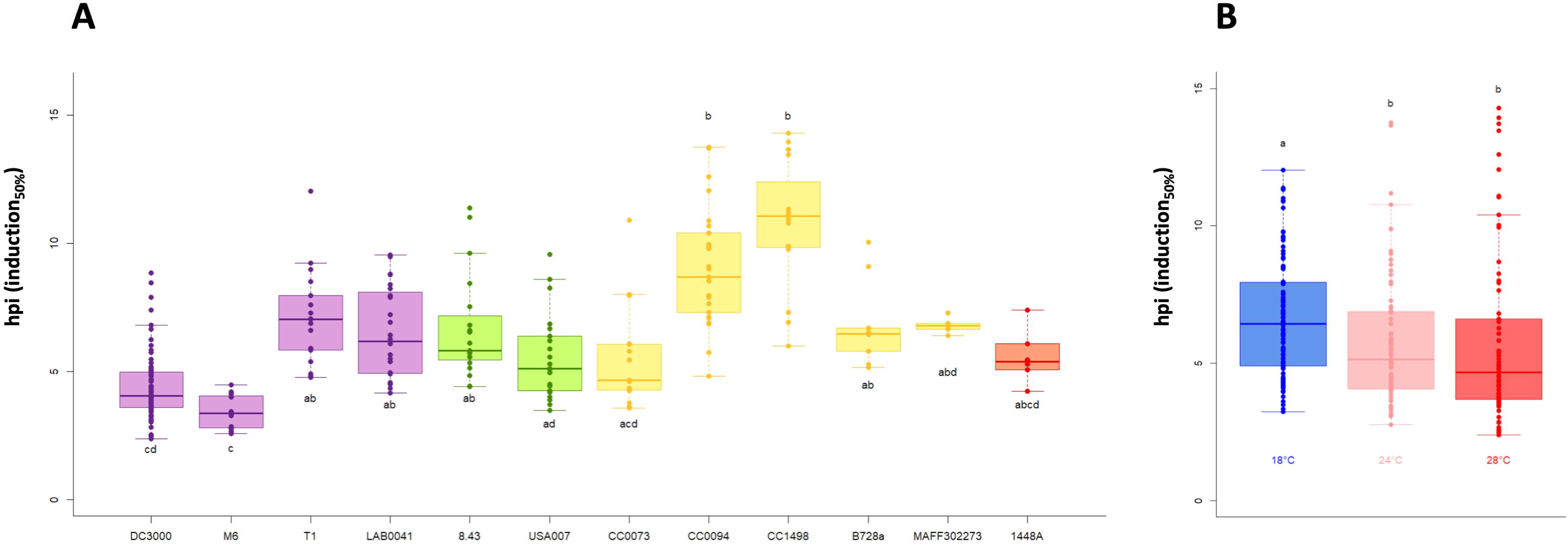
Estimation of velocity of HR induction. The proxy for velocity was calculated as the time point when half of conductivity increase is reached for each *avrB*-expressing strain inducing HR at least at one temperature condition. (a) Strains are sorted by phylogroup from left to right, PG1a (purple), PG1b (green), PG2d (yellow), PG3a (red). Boxes were calculated from 5 up to 73 replicated experiments per strain (single dots). Boxes associated with different letters represent significantly different means (p < 0.05, according to a Kruskal-Wallis’s test followed by post-hoc Dunn’s test). (b) Boxes were calculated from 70 to 97 replicated experiments per temperature (single dots). Boxes associated with different letters represent significantly different means (p < 0.05, according to a Kruskal-Wallis’s test followed by post-hoc Dunn’s test).

## Discussion

### New perspectives on virulence thermoregulation processes

The diverse behaviors of T3SS efficiency observed in relation to incubation temperature indicate that *P. syringae* has a very diverse adaptive potential for causing disease in the face of various temperatures. This observation is another piece of evidence out of the many reported over the past 30 years regarding the role of temperature in virulence regulation in plant-associated bacteria (reviewed in references Smirnova et al., 2001; Velásquez et al., 2018). Our work has presented new examples of relatively warm-temperature dependency for virulence efficiency, consistent with previous findings about gene expression beyond and within the *P. syringae* species complex (Czajkowski et al., 2017; Huot et al., 2017; Krishna et al., 2022; Peñaloza-Vázquez et al., 1997; Wei et al., 2008; Wei et al., 2010). Altogether, these pieces of evidence contradict the long-standing notions that, firstly, “almost all virulence genes of plant pathogenic bacteria […] exhibit increased transcription at temperatures well-below the respective growth optima” (Smirnova et al., 2001) and, secondly, that “the increase of bacterial growth in plants at elevated temperature cannot be explained by variations in intrinsic bacterial growth rate or secretion of effector proteins at different temperatures” (Wang et al., 2009). Our findings support a distinctly different perspective that aligns with more recent articles that present an updated framework (Huot et al., 2017; Tribelli and López, 2022; Velásquez et al., 2018).

### Effector repertoire contribution in HR induction

Although *avrB* expression was assessed for all the strains, two of them (J35 *avrB* and MAFF302273 *avrB*) were categorized as non-inducers for at least two of the tested temperatures. The lack of HR induction could result from T3SS inactivation in *A. thaliana*, potentially because of a quick PTI onset (Crabill et al., 2010). On the other hand, it could also be a matter of speed of ETI onset, whose rapidity prevents cell death occurrence (Künstler et al., 2016). Otherwise, it may be due to the presence in the repertoire of the strain of some suppressive effector(s) (Jamir et al., 2004), or some effector(s) targeting a molecular actor involved in both PTI and ETI, or simply blocking the PTI – required in the first place for ETI activation (Ngou et al., 2021). Although our analysis of effector repertoire composition (PsyTEC; Laflamme et al., 2020) did not allow to identify one single or group of effectors antagonistic to AvrB, we cannot rule out the occurrence of ‘meta-effector’ interactions (Kubori et al., 2010; Martel et al., 2022). Overall, the in-depth-study of the effector repertoires of all the strains used in our work could help to explain the differences observed in terms of HR intensity, with one or more effectors that may inhibit HR but also enhance it conversely – as observed for strain CC1498 *avrB* versus strain CC0094 *avrB*, for example.

### Temperature and plant immunity

Using HR induction as a proxy for T3SS efficiency, we implicitly accounted for plant defense mechanisms that play a role in the response (Künstler et al., 2016). In *A. thaliana*, the role of temperature in plant immunity is more complex than reported by previous studies. In 2009, Wang and colleagues concluded that a mild increase in temperature impaired both PTI and ETI, suggesting a general deficiency in the plant defense system at elevated temperatures (28-30°C) (Wang et al., 2009). Subsequently, a straightforward dichotomy was established: less efficient ETI (effective from 10 to 23°C, with a peak at 18°C) versus more efficient PTI (effective from 23 to 32°C, with a peak at 28°C) as the temperature increases (Cheng et al., 2013). This model was later validated by the demonstration that, since ETI relies on distinct processes (*i.e.*, virulence suppression and programmed cell death/HR), only HR was inhibited by elevated temperatures as the virulence suppression was maintained (Menna et al., 2015). However, despite their demonstration that ion leakage from *A. thaliana* leaf disks infiltrated with the avirulent DC3000 strain is completely inhibited at 28°C-30°C, our results for this *avrB*-expressing strain and CC0094 *qvrB* and CC1498 a*vrB* contradict this finding.

Temperature variations may affect the kinetics of effector translocation and/or binding to their targets and hence impact the speed and efficiency of signaling cascades involved in the defense response. Comparison of the time point when half of the final conductivity measured at 24 hpi – hpi_50%_ – among temperatures for a given HR-inducer strain revealed that the elevation of the temperature led to an acceleration of electrolyte leakage as temperature increased. As this observation was maintained across the strains that induced HR at different temperatures, it is tempting to associate this general effect of accelerated response with the effect of the higher temperature on the plant, and not on the T3SS activation or efficiency. Nevertheless, across all strains, we demonstrated that some were faster than others to induce HR, also indicating some pathosystem-specific behaviors.

### Temperature and effector secretion properties

Interestingly, a few examples of temperature-dependency for T3E secretion have been shown to be ‘protein/effector specific’ – mentioning some specific secretion properties of AvrB (Van Dijk et al., 1999). Likewise, the two avirulent DC3000 strains with which HR suppression at elevated temperatures was demonstrated carried either the HopZ1a or AvrRpt2 effector (Menna et al., 2015). HopZ1a, AvrRpt2 and AvrB are recognized by the NLRs ZAR1, RPS2 and RPM1, respectively, which are all members of the subgroup of CC-NLRs. The unclear variability of temperature-sensitivity across different NLR subgroups (CC-NLRs versus TIR-NLRs) (Hua, 2014; Menna et al., 2015) suggests that sensitivity may not be conserved in all NLRs or CC-NLRs, given their distinct structural and functional properties. In this light, it is reasonable to attribute the differences observed between our work and the one of Menna and colleagues (Menna et al., 2015) to the different ‘ligand-receptor’ couples used. Echoing the differences in ‘protein/effector specific’ secretion efficiency, there are likely to be differences in temperature-sensitivity for effector recognition by R proteins, providing strong support to our choice to introduce the same effector in all strains to ensure a proper comparison of HR induction dynamics. Overall, we assume that the impact of temperature on T3SS-secretion/ETI/HR-triggering efficiencies can be highly pathosystem-specific.

### Virulence thermoregulation in relation to disease development

Since various virulence factors may be controlled by the same thermoregulation components (Klinkert and Narberhaus, 2009; Smirnova et al., 2001), one intriguing question is whether the effect of temperature on virulence factors is consistent (i) for a given strain, and (ii) with temperature conditions at which disease occurrence is generally reported. In other words, do our results correspond to what is known about disease epidemiology under field conditions? For *P. syringae* pv. *phaseolicola*, low-temperature-dependency was established for phaseolotoxin production and T6SS regulation, in line with the development of halo blight of bean favored at temperatures below 25°C (Goss, 1940; Nüske and Fritsche, 1989). However, T3SS-related gene expression inhibition has been reported at 18°C (Arvizu-Gómez et al., 2013), and our results indicate that the strain 1448A is thermo-insensitive in its ability to induce HR. Regarding *P. syringae* pv. *actinidiae*, our results confirmed a low-temperature dependency for T3SS efficiency (Puttilli et al., 2022), which is entirely in line with the environmental conditions – in late winter or early spring – under which *P. syringae* pv. *actinidiae* is reported to cause disease or damage in the kiwifruit orchards (Donati et al., 2020; Kim et al., 2017). On the other hand, concerning strains CC0073 and CC0094, isolated from cantaloupe during blight epidemics in southern France in the 90s, the authors have reported that the climatic conditions that favored disease outbreak and development were heavy rainfall associated with a mean minimum temperature below 12-13°C (under which the physiology of cantaloupe comes to a standstill) in 95% of cases, whereas our results demonstrated a major T3SS efficiency at higher temperatures (Mention et al., 2004; Morris and Pitrat, 1998; Riffaud, 2002). Finally, strain DC3000 displays a consistent behavior, exhibiting both unaltered T3SS efficiency and coronatine production in response to temperature, at least within the 18°C to 28°C range (Weingart et al., 2004). Its consistency in response to various environmental parameters, associated with high reproducibility of data related to it, has certainly contributed to making it the model strain it has become. However, this strain was generated in the laboratory as a rifampicin-resistant derivative of the strain DC52, which originated from Guernsey in 1960 (Cuppels, 1986; Xin et al., 2018). Therefore, no conclusive connection could be established between its ecology and its virulence behavior in response to temperature, as this model strain has evolved for many years in laboratories, far from a field context.

Our results also provide evidence that growth optima – at least *in vitro* – are not correlated with T3SS efficiency. This contributes to the debate about the question of whether the temperature conditions that favor virulence (specifically T3SS) are identical to those reported as favorable for disease development. Answering this question is challenging due to both supporting and opposing examples when accounting for the various traits characterizing phytopathogenic bacteria that are thermo-sensitive, and each pathogen is characterized by growth and virulence mechanisms that are activated at precise temperatures, generating a complex scenario in which temperature can alter the outcome of the interaction between plant and bacterium by differentially influencing a spectrum of virulence traits (Tribelli and López, 2022). Consequently, it is likely to be very difficult to simplify or generalize the behavior of different strains relative to the influence of temperature on their virulence.

### Inter-strain variability

We observed that the behavior of high temperature-preference/dependency is exclusive to strains from phylogroup 2d, whereas the opposite case is specific to strains belonging to phylogroups 1a and 1b. This suggests that temperature responsiveness patterns might vary among strains but remain consistent within the same phylogroup. While the temperature sensitivity feature is widely distributed among the strains, the preference/dependency on low or high temperatures in case of thermosensitivity is conserved within a same phylogroup. Therefore, the preference/dependency on low or high temperatures could be linked to the evolutionary history of pathogens, which may have been shaped by temperature-related selection pressures. Investigating this question would help shed light on the origin of the phenomenon. On the other hand, one important conclusion of our work is the significant variability in behavior among strains, even when closely related at the genomic level (as illustrated with the 3 ‘CC’ strain-quasi-clones). Over the years of research on these issues, examples of such inherent variability in terms of virulence have been largely overlooked, with only a few cases reported in literature, both beyond and within the *P. syringae* species complex (Baron et al., 2001; Hasegawa et al., 2005; Rohde et al., 1998; Weingart et al., 2004). Here, we emphasize the urgent need to study the diversity among strains, since evidence of their inherent variability has been documented. Although investigating model strains has facilitated the understanding and characterization of intricate molecular mechanisms in plant-pathogen interactions, it is now imperative to consider the full spectrum of *P. syringae* species complex diversity (Berge et al., 2014; Gomila et al., 2017), both within and outside of the agronomic context.

## Experimental procedures

### Bacterial strains, culture conditions and transformation

Thirteen *Pseudomonas syringae* strains were selected for this study (Supplementary Table S1; Supplementary information). Wild-type (WT) *P. syringae* strains were grown at 28°C on King’s B agar (KB) (King et al., 1954) – supplemented with rifampicin at 50µg/mL for DC3000 and B728a strains. Kanamycin at 50µg/mL was added for the culture of *avrB*-expressing strains. Single colonies of each strain were picked and grown overnight in liquid KB medium – supplemented with rifampicin at 50µg/mL for DC3000 and B728a strains and with kanamycin at 50µg/mL for *avrB*-expressing strains, with agitation at 190 rpm at 28°C.

Selected strains were transformed with the pure plasmid carrying *avrB* gene whose sequence was originally isolated from the soybean pathogen *Pseudomonas syringae* pv. *glycinea* (pVB01 Km^R^; Innes et al., 1993) through electroporation process. Briefly, fresh suspensions of WT strains grown overnight in liquid KB (190 rpm, 28°C) were washed three times and resuspended in 50-100uL of cold 10% glycerol. Pure plasmid (50-250ng) was added to the suspensions which were then transferred into cold 2mm electroporation cuvettes (BioRad, CA, USA) Electroporation was performed with a MicroPulser^TM^ machine (BioRad, CA, USA) set at 1.8-2.5kV. Cells were immediately recovered into fresh nonselective liquid KB medium and incubated at 28°C for 1-2h at 190 rpm. Transformed strains were selected using kanamycin at 50µg/mL (plus rifampicin at 50µg/mL for DC3000 and B728a strains) and the presence of the plasmid was confirmed by colony PCR using Promega GoTaq™ DNA polymerase (Fisher Scientific, USA) and specific primers (FOR: ATGGGCTGCGTCTCGTCAAAAAGCA, REV: TTAAAAGCAATCAGAATCTAGCAAG, annealing temperature = 65°C).

### *avrB* expression control

Relative expression levels of *avrB* in transformed strains were evaluated by real-time PCR. Four mL of fresh suspension of *avrB*-transformed *P. syringae* strains grown overnight in selective medium were centrifuged (5min at 4500g at RT) and total RNA was extracted from pellets using TRIzol substitute (EuroGold TriFast^TM^ Kit, EuroClone, Italy) according to manufacturer instructions. Extraction yield and quality of the transcripts were checked by spectrophotometry (NanoDrop^TM^, Thermo Fisher Scientific, USA). Samples with absorbance ratios A_260nm_/A_230nm_ and/or A_260nm_ /A_280nm_ outside the range of [2 ± 0.5] were subjected to a lithium chloride treatment to increase their purity (2.5M LiCl, 50mM EDTA, pH 8.0). One to two µg of RNA per sample underwent DNase treatment (TURBO^TM^ DNase, Life Technologies, USA) then the single-stranded complementary DNA synthesis was performed from 0.5-1µg of template RNA using SuperScript^TM^ III Reverse Transcriptase enzyme (Invitrogen, USA) according to manufacturer instructions. Samples get 10-fold diluted and the procedure was performed using Luna® Universal qPCR Master Mix (New England Biolabs, USA). To normalize the expression of the target gene, *rpoD* was chosen as a stable and robust housekeeping gene in *Pseudomonas* (Narusaka et al., 2011; Savli et al., 2003). Amplification was run on a QuantStudio^TM^ 3 real-time PCR system (Applied Biosystems, USA) (primers *avrB* FOR: ATACTCTGCATTCGCCTCCG, REV: TCAACGACTCTGGCAAGGAC; primers *rpoD* FOR: AACTTGCGTCTGGTGATCTC, REV: ATCAAGCCGATGTTGCCTTC). Expression of *avrB* was determined using the 2^−ΔΔCt^ method and normalized against *rpoD*, and fold-change for each strain was established relative to the strain DC3000 *avrB*.

### Ion-leakage experiments

Hypersensitive response (HR) induction was assessed by quantifying programmed cell death through ion-leakage measurement, as previously published (Imanifard et al., 2018; Jayaraman et al., 2021; Johansson et al., 2015) with slight modifications provided as Supplementary Material.

### Statistical analysis

For each technical replicate, the conductivity value from the first measure was subtracted to values measured at each successive time points for normalization (Puttilli et al., 2022). For each strain-temperature modality within each replicated experiment, conductivity was calculated as the average of the values of the three technical replicates for each time point, and ion-leakage curves over time were established accordingly. Area under each single conductivity progress curve was calculated (AUCPC), as well as variance and mean AUCPC per modality. Moreover, conductivity values corresponding to 24 hpi as well as the time point for which half of this value was reached for each experiment were calculated via linear interpolation, and this variable hpi_50%_ was used as a dynamic parameter, indicating how fast a given strain induced HR at a given temperature. Comparison of strains based on this parameter was assessed by variance analysis (Kruskal-Wallis’s test) followed by a post-hoc Dunn’s test. The same procedure was conducted to compare the three different temperatures regardless of the strain. Data were represented by boxplots.

Each single conductivity progress curve constituted from 6 to 9 time points and was considered as one statistical individual (*i.e.*, one iteration) to fit Linear Mixed-effects Models (LMM). Details of the approach are provided as Supplementary Material.

### *In vitro* HrpZ secretion

Twenty mL of fresh suspension of *avrB*-expressing *P. syringae* strains grown overnight in selective medium were centrifuged (5min at 4500g at room temperature) and pellets were washed three times in 10mL of sterile minimal medium (HIM). OD_600_ was measured with a spectrophotometer (Evolution™, ThermoFisher Scientific, USA) and bacterial cells were resuspended in sterile HIM to 7 x 10^8^ CFU/mL (OD_600nm_ = 0.7; final volume = 150mL). Suspensions were divided into three portions of 50mL, each was separately incubated at 18°C, 24°C or 28°C with agitation at 190 rpm. After 5 and 24 hours of incubation, 25mL of each bacterial suspension were sampled and OD_600_ was measured again. Cell-free culture supernatant was obtained by centrifugation and filtration, and secreted proteins were precipitated with trichloroacetic acid (TCA) 5% as previously described (Flaugnatti and Journet, 2017; Kobayashi et al., 1997).

For immunoblotting, proteins were separated on SDS–polyacrylamide gel electrophoresis and then transferred onto a nitrocellulose blotting membrane (GE HealthCare, USA) using an electrophoretic apparatus. Protein detection was performed using anti-HrpZ (1:5000) (Alfano et al., 1996) and anti-rabbit (1:10000) conjugated with horseradish peroxidase as secondary antibody. To confirm the correct loading ratio among wells, one gel was stained with a Coomassie R-250 staining solution (25% isoprophyl alcohol, 10% acetid acid, 0.05% Coomassie R-250) and destained with 10% acetic acid.

### *In vitro* growth measurement

Four mL of fresh suspension of *avrB*-expressing *P. syringae* strains grown overnight in selective medium were centrifuged (5min at 4500g at room temperature) and pellets were washed three times in 1mL of fresh liquid KB. Optical density at 600nm was measured by spectrophotometer (Evolution™, ThermoFisher Scientific, USA) and bacterial cells were then resuspended to reach 2 x 10^7^ CFU/mL (OD_600nm_ = 0.02; final volume = 2mL). Suspensions were aliquoted in a microtiter plate (200µL per well), with 6 technical replicates per strain, and plates were incubated at 18°C, 24°C and 28°C. Liquid KB (supplemented or not with the antibiotics) was used as negative control. Absorbance at 600nm was measured every 1.5-2 hours during the first 8 hours of the experiment, and then two or three times between 18 and 24 hpi, obtaining from 6 to 9 absorbance measures per well. To eliminate unspecific absorbance signals due to the color of KB medium and the antibiotics, the value from the first measure was subtracted from values measured at each successive time points. Absorbance was then calculated as the average of the values of the six technical replicates for each time point, and bacterial growth curves over time were established accordingly. Each strain-temperature modality was reproduced at least twice, except for T1.

## Acknowledgments

We thank Pr. Boris A. Vinatzer, Dr. Perrine Portier, and Dr. Marco Scortichini for kindly providing *P. syringae* strains used in this study in addition to our own strains. We also thank Pr. Sheng-Yang He for providing the antibody anti-HrpZ. Special thanks to Pr. Roberto Chignola and Dr. Davide Martinetti for their help in statistical analyzes of ion-leakage data. Finally, we also thank the members of the Biotechnology Department of the University of Verona who provided technical support in conductivity measurements.

## Data Availability Statement

All data needed to evaluate the conclusions in the paper are present in the paper and/or the Supporting Information. Additional data related to this paper may be requested to the authors.

## Supplementary figure legends

**Figure S1. Variance of the mean area under conductivity progress curve (AUCPC) values obtained for each 39 modalities *avrB*-expressing strain x temperature.** Number of replicated experiments for each strain varied from 3 up to 27. Linear regression trend curve was fitted to the data, with y=323.89x – 150932 and R²=0.8588, showing a positive correlation between mean AUCPC and their variance (Spearman’s coefficient = 0.91; p < 0.05).

**Figure S2. avrB expression among the mutant strains.** RNA was extracted from bacterial suspensions grown overnight at 28°C in liquid KB medium supplemented with kanamycin (50μg/mL) and rifampicin (50μg/mL, for DC3000 *avrB* and B728a *avrB* only). Real-time PCR was performed using *rpoD* as the housekeeping gene. Expression levels were established using the 2–ΔΔCt method using DC3000 *avrB* as a reference (fold change = 1). Error bars represent standard error. Data represent the mean of three independent biological replicates, each repeated at least twice.

**Figure S3. *In-vitro* growth curves for DC3000 avrB (a), USA007 avrB (b) and CC0094 avrB (c).** Bacterial growth was measured for 24 hours at 18°C (blue), 24°C (pink) and 28°C (red). Overnight grown bacteria were resuspended in liquid KB medium supplemented with kanamycin (50μg/mL) and rifampicin (50μg/mL, for DC3000 *avrB* only) at an initial load of 107 CFU/mL (optical density measured at 600nm = 0.01). Data represent one single representative biological replicate with all strains and temperature conditions evaluated simultaneously with six technical replicates each. Error bars represent standard error.

**Figure S4. Electrolyte leakage curves over time obtained post-infiltration for CRA-FRU 8.43 in *Actinidia arguta* (a) and CC0094 *in A. thaliana* Col-0 (b).** Infiltrated leaf disks were incubated at 18°C (blue) and 28°C (red). Conductivity following infiltration of the mock treatment (10mM MgCl_2_) and incubation at 18°C is represented by the black dotted line. Data represent one single representative biological replicate with all strains and temperature conditions evaluated simultaneously with three technical replicates each. Error bars represent standard error.

