## Supporting information for "Some like it hot: efficiency of the type III secretion system has multiple thermosensitive behaviors in the *Pseudomonas* syringae complex"

Emma Caullireau^1,2^, Davide Danzi^1^, Vittoria M. Tempo^1^, Mattia Pandolfo^1^, Cindy E. Morris^2^, Elodie Vandelle^1^*

^1^ Department of Biotechnology, University of Verona, 37134 Verona, Italy.

^2^ UR0407 Plant Pathology, INRAE, 84140 Montfavet, France.

*Elodie Vandelle

**This PDF file includes:**

Supplementary Material and Methods

Figures S1 to S4

Tables S1 to S3

Supplementary Material and Methods

Bacterial strain selection

Selected strains represented four phylogroups/clades (1a, 1b, 2d, and 3a). The geographical origin and isolation source of the strains varied, encompassing annual crops (*e.g.* M6, T1) (Debener et al., 1991; Whalen et al., 1991), woody plants (*e.g.* J35, CRA-FRU 8.43, MAFF302273) (Ferrante and Scortichini, 2010; Sawada et al., 1999; Takikawa et al., 1989), and environmental reservoirs like rivers or snow (*e.g.* LAB0041, USA007) (Berge et al., 2014; Morris et al., 2010). Genomic relatedness within a phylogroup was also considered, aiming for a balance of very divergent and closely related strains, such as the three ‘CC’ strains (CC0073, CC0094, and CC1498) that are quasi-clones (ANI > 99.83%) (Monteil et al., 2016; Morris et al., 2000; Morris et al., 2008). Model strains DC3000, B728a and 1448A were also included in the set as there is a plethora of information about their biology (Cuppels, 1986; Loper and Lindow, 1987; Taylor et al., 1996).

Ion-leakage experiments

Strains expressing *AvrB* were streaked from glycerol stocks on King’s B agar (KB) (King et al., 1954) – supplemented with kanamycin at 50ug/mL and rifampicin at 50µg/mL for DC3000 and B728a strains. After 48h of incubation at 28°C, single colonies of each strain were picked and grown overnight in liquid selective KB medium, with agitation at 190 rpm at 28°C. Similar procedure was followed for CC0094 WT (grown in nonselective KB). *A. thaliana* Col-0 plants were cultivated in a mixture of peat soil and perlite (4/1, v/v) in a growth chamber set at 24°C/21.5°C thermoperiod and 8 h light–16 h dark photoperiod with 70% of relative humidity for 6 to 8 weeks. Three healthy plants with six 5-mm leaf disks detached from a same plant with a cork borer were used for each experimental condition. To recognize the plant they were detached from, leaf disks were marked with a colored felt-tip and were then transferred into a 50mL tube containing 10mM MgCl_2_ solution during the time of bacterial suspension preparation. Ten mL of fresh bacterial suspensions were centrifuged (5min at 4500g at room temperature) and pellets were washed three times in 10mL of sterile 10mM MgCl_2_ solution. Optical density at 600nm was measured by spectrophotometer (Evolution™, ThermoFisher Scientific, USA) and bacterial cells were finally resuspended in sterile 10mM MgCl_2_ solution to reach 10^8^ CFU/mL (OD_600nm_ = 0.1; final volume = 30mL). Leaf disks were infiltrated with the bacterial suspensions using vacuum (applied using a 50mL syringe or a vacuum chamber). Sterile 10mM MgCl_2_ solution or *hrp*-inducing medium (HIM; using 27mM glycerol as carbon source; pH 5.5) (Huynh et al., 1989) were used as negative controls. Infiltrated leaf disks were rinsed twice for 15m in 40mL of milliQ water under gentle agitation (90 rpm) at room temperature to eliminate suspension/mock solution trace and ions leaked due to mechanical damage. Infiltrated leaf disks were then distributed into 12-well plate containing 2mL of milliQ water per well in order to have: (i) six disks per well, (ii) two disks from each of the three different plants per well, (iii) three identical wells (one plate column) per each experimental modality. Initial conductivity values (µS.cm^-1^) in each well was measured prior plates incubation (120 rpm; constant light of 80 μmol.m^-2^.s^-1^) at different temperatures (18°C, 24°C and 28°C) using a compact conductometer (LAQUAtwin EC-11, HORIBA Scientific, Japan). Conductivity increase was measured every 1.5-2 hours during the first 8 hours of the experiment, and then twice between 18- and 24-hours post-inoculation (hpi), obtaining from 6 to 9 conductivity measures on average per well. Each experimental modality (*i.e.* one specific *avrB*-expressing strain infiltrated and then incubated at a certain temperature) was reproduced at least three times (at least 3 `biological replicates’ or `replicated experiments’).

To assess the hypersensitive response induction in kiwifruit, adult plants of *Actinidia arguta* var. Kens Red were recovered from a local nursery (Vivai Guardini, Pescantina, Verona, Italy) and maintained in pots in a greenhouse following natural environmental conditions. Strains CRA-FRU 8.43 WT and *avrB* were streaked from glycerol stocks on LB agar and incubated at 22°C for 48h. Cells were recovered from this plate and spread on fresh LB agar plates and incubated overnight at 22°C. Cells were finally scraped from plates and suspended in a 10 mM MgCl_2_ solution to reach 10^9^ CFU/mL (OD_600nm_ = 1). Leaf disks (8 mm) were made from young leaves of a single *A. arguta* var. Kens Red plant and vacuum infiltrated using a syringe following the same protocol used for *A. thaliana*.

Effector repertoire characterization

The effector repertoires of ten *P. syringae* strains from our set were already publicly available (Dillon et al., 2019; Laflamme et al., 2020). We characterized the effector repertoires of the remaining strains (LAB0041, CC0073, CC1498) from their draft genomes (provided by Dr. Cindy Morris). The genomes were analyzed following a custom bioinformatic pipeline created with the help of the Nextflow framework (Di Tommaso et al., 2017). The pipeline was composed of several steps: i) the FastANI tool was used for rapid many-to-many ANI calculation (Jain et al., 2018); ii) BUSCO was used to assess both the genome quality and protein annotations quality (Manni et al., 2021); iii) the Prokka pipeline was used for the protein annotation (Seemann, 2014). The three genome protein annotations produced were aligned to the amino acid sequences of 529 T3SE proteins (obtained from the *Pseudomonas syringae* Type III Effector Compendium, PsyTEC) (Laflamme et al., 2020), list available at: <https://guttman.csb.utoronto.ca/files/2023/07/PsyTEC_Public_Resource.xlsx>) in a reciprocal best-hit (RBH) alignment via BLASTP, run with an e-value threshold of 1e-24 (Altschul et al., 1997; Daubin et al., 2002; Eisen, 2000; Hernández-Salmerón and Moreno-Hagelsieb, 2020). The alignment outputs were further refined with an R script, which kept, for each genome, only the unique proteins present in the T3SE list with a minimum RBH alignment coverage of 60% and a minimum identity score of 95%. These resulting proteins were considered as candidate effectors for the relative genome and were used to produce a presence/absence effector table for the 3 genomes, which was then merged with the existing one for the 10 previously characterized strains. Information was synthetized to present family-level classified effector repertoires.

Statistical analysis (Linear Mixed-effects Models)

This allowed us to account for the important variability among the diverse experimental replicates of a same modality and, most importantly, to consider the entire time course of electrolyte leakage, rather than focusing solely on isolated time points. The mathematical representation of LMM is: *Y = Xβ+ Zʋ+ ε* ; where *Y* represents the dependent variable (the measured feature, *i.e.* conductivity increase), *Xβ* stands for the fixed effects (*i.e.* the potential explanatory variables we want to assess the effect on *Y* ), *Zʋ* stands for the random effects (all the unknown parameters that may be responsible for the huge variability among the different replicated experiments and which we are not directly interested to), and *ε* represents the residual. Using LMM in RStudio, we evaluated (i) the strain effect (*i.e.* if the ion-leakage following *avrB*-expressing strain infiltration was significantly different from the ion-leakage due to mock treatment infiltration) to categorize each strain as ‘HR-inducer’ or ‘non-inducer’ at each of the tested temperatures; and (ii) the temperature effect (*i.e.* if the ion-leakage following *avrB*-expressing strain infiltration was significantly different as the incubation temperature changed) on the strains considered as HR-inducers at least at two of the tested temperatures. In this way we estimated the values of model parameters and the associated p-values.

**Supplementary Figures**

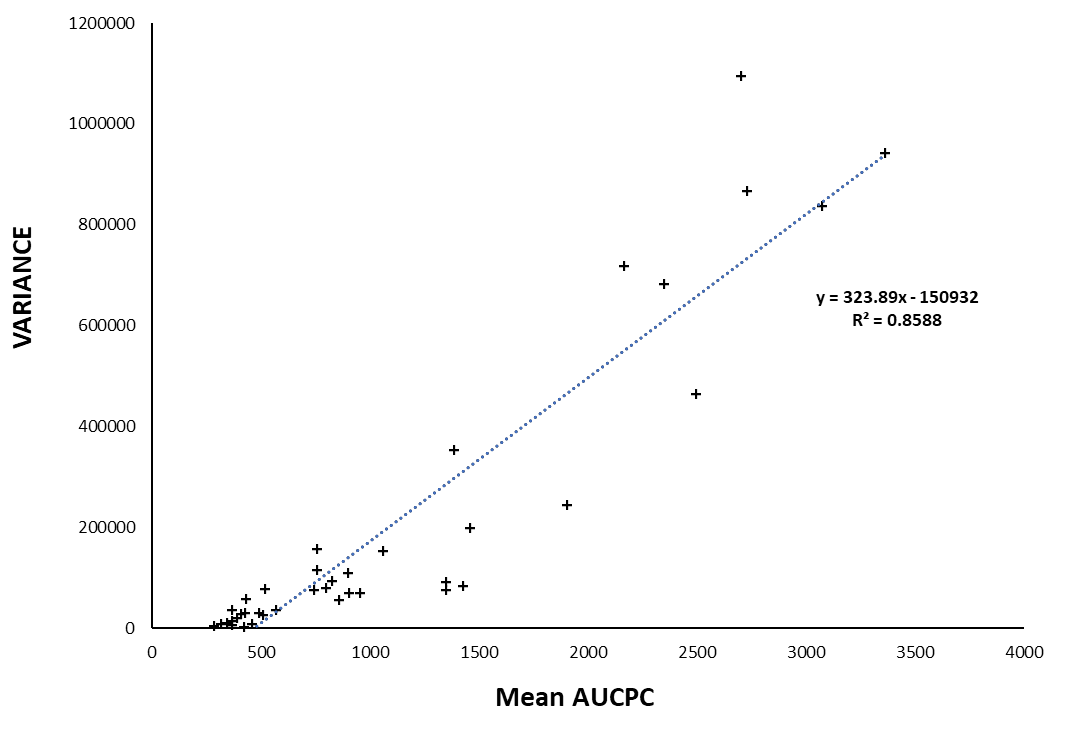

Figure S1. Variance of the mean area under conductivity progress curve (AUCPC) values obtained for each 39 modalities *avrB*-expressing strain x temperature. Number of replicated experiments for each strain varied from 3 up to 27. Linear regression trend curve was fitted to the data, with y=323.89x – 150932 and R²=0.8588, showing a positive correlation between mean AUCPC and their variance (Spearman’s coefficient = 0.91; p < 0.05).

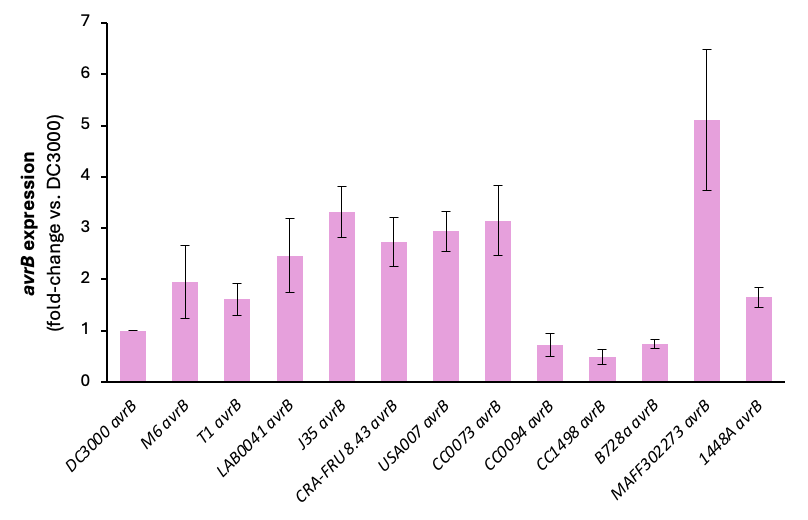

Figure S2. *avrB* expression among the mutant strains. RNA was extracted from bacterial suspensions grown overnight at 28°C in liquid KB medium supplemented with kanamycin (50μg/mL) and rifampicin (50μg/mL, for DC3000 *avrB* and B728a *avrB* only). Real-time PCR was performed using *rpoD* as the housekeeping gene. Expression levels were established using the 2^–ΔΔCt^ method using DC3000 *avrB* as a reference (fold change = 1). Error bars represent standard error. Data represent the mean of three independent biological replicates, each repeated at least twice.

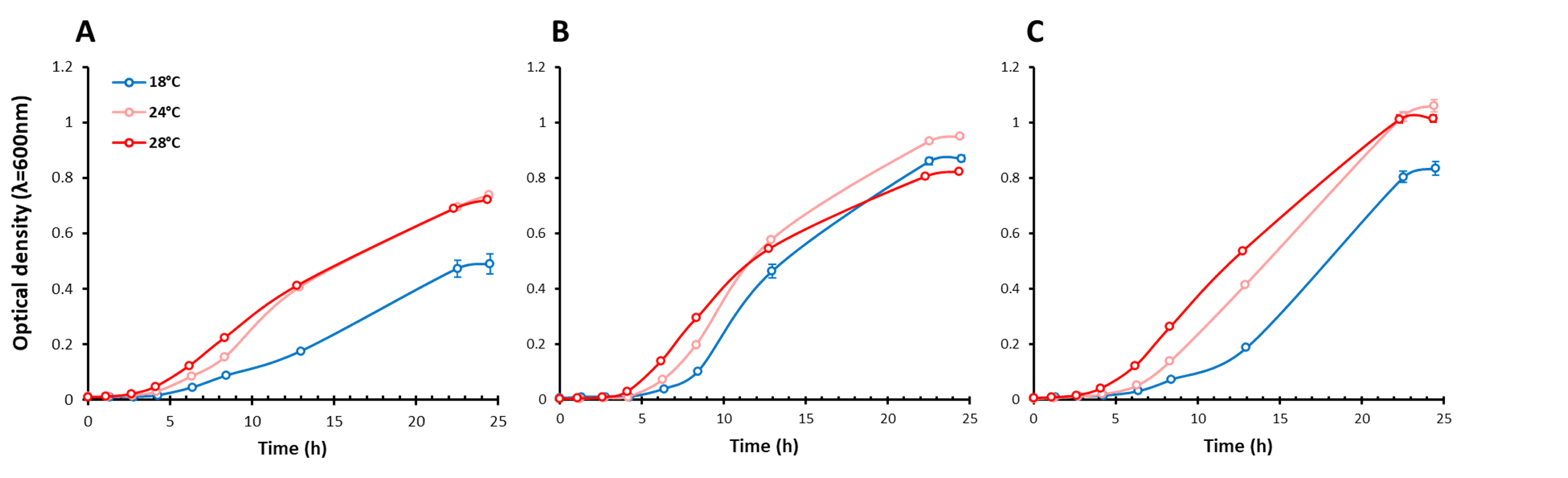

**a**

**b**

**c**

Figure S3. *In-vitro* growth curves for DC3000 *avrB* (a), USA007 *avrB* (b) and CC0094 *avrB* (c). Bacterial growth was measured for 24 hours at 18°C (blue), 24°C (pink) and 28°C (red). Overnight grown bacteria were resuspended in liquid KB medium supplemented with kanamycin (50μg/mL) and rifampicin (50μg/mL, for DC3000 *avrB* only) at an initial load of 10^7^ CFU/mL (optical density measured at 600nm = 0.01). Data represent one single representative biological replicate with all strains and temperature conditions evaluated simultaneously with six technical replicates each. Error bars represent standard error.

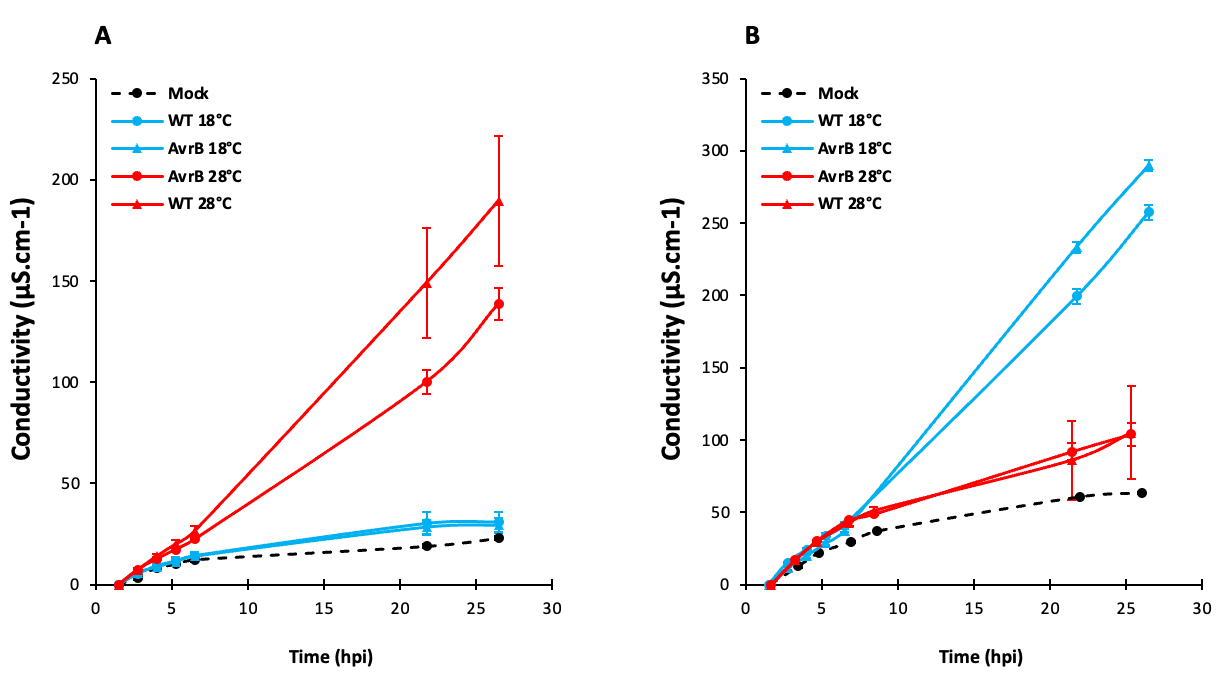

**b**

**a**

Figure S4. Electrolyte leakage curves over time obtained post-infiltration for CRA-FRU 8.43 in *Actinidia* *arguta* (a) and CC0094 in *A. thaliana* Col-0 (b). Infiltrated leaf disks were incubated at 18°C (blue) and 28°C (red). Conductivity following infiltration of the mock treatment (10mM MgCl_2_) and incubation at 18°C is represented by the black dotted line. Data represent one single representative biological replicate with all strains and temperature conditions evaluated simultaneously with three technical replicates each. Error bars represent standard error.

**Supplementary Tables**

**Supplementary Table S1. Set of selected *P. syringae* strains.**

| **PG** | **STRAIN** | **FULL NAME** | **ALIAS IN COLLECTIONS** | **ISOLATION SOURCE** | **COUNTRY** | **YEAR** | **REFERENCE** |
| --- | --- | --- | --- | --- | --- | --- | --- |
| 1a | DC3000 | *P*. *syringae* pv. *tomato* | CFBP 7438, ICMP 18429 | *Solanum lycopersicum* | United-Kingdom | 1960 | (Cuppels, 1986) |
|  | M6 | *P*. *syringae* pv. *maculicola* | CFBP 4314, NCPPB 1766, LMG 5560 | *Brassica oleracea* var. Botrytis | United-Kingdom | 1965 | (Debener et al., 1991) |
|  | T1 | *P*. *syringae* pv. *tomato* | - | *Solanum lycopersicum* | Canada | 1986 | (Whalen et al., 1991) |
|  | LAB0041 | *P*. *syringae* | - | Epilithic biofilm | France | 2009 | (Berge et al., 2014) |
| 1b | J35 | *P*. *syringae* pv. *actinidiae* | CFBP 4909, ICMP 9617, NCPPB 3739 | *Actinidia deliciosa* cv. Hayward | Japan | 1984 | (Takikawa et al., 1989) |
|  | CRA-FRU 8.43 | *P*. *syringae* pv. *actinidiae* | - | *Actinidia chinensis* cv. Hort16 | Italy | 2008 | (Ferrante and Scortichini, 2010) |
|  | USA007 | *P*. *syringae* | CFBP 8574 | Creek water | United States of America | 2007 | (Morris et al., 2010) |
| 2d | CC0073 | *P. syringae* pv. *aptata* | CFBP 5430 | *Cucumis melo* | France | 1997 | (Morris et al., 2000) |
|  | CC0094 | *P*. *syringae* pv. *aptata* | CFBP 8529 | *Cucumis melo* | France | 1997 | (Morris et al., 2000) |
|  | CC1498 | *P*. *syringae* | - | Snowfall | France | 2006 | (Morris et al., 2008) |
|  | B728a | *P*. *syringae* pv. *syringae* | CFBP 8502, ICMP 18427, NCPPB 4487, LMG 26717 | *Phaseolus vulgaris* | United States of America | 1987 | (Loper and Lindow, 1987) |
|  | MAFF302273 | *P*. *syringae* pv. *aceris* | CFBP 2339, ICMP 2802, NCPPB 958, LMG 2106 | *Acer* sp. | United States of America | 1939 | (Sawada et al., 1999) |
| 3a | 1448A | *P*. *syringae* pv. *phaseolicola* | CFBP 7087, NCPPB 4478 | *Phaseolus vulgaris* | Ethiopia | 1985 | (Taylor et al., 1996) |

**Supplementary Table S2. Matrix of effectors found among the *P. syringae* strains.** Effector repertoires of all strains used in this study were already available publicly, with the exception of CC0073, CC1498 and LAB0041 whose repertoires were characterized following the same procedure, as described (Laflamme et al., 2020).

| olonne1 | **DC3000** | **M6** | **T1** | **LAB0041** | **J35** | **CRA-FRU 8.43** | **USA007** | **CC0073** | **CC0094** | **CC1498** | **B728a** | **MAFF302273** | **1448A** |
| --- | --- | --- | --- | --- | --- | --- | --- | --- | --- | --- | --- | --- | --- |
| AvrA | 0 | 0 | 1 | 0 | 0 | 0 | 0 | 0 | 0 | 0 | 0 | 0 | 0 |
| AvrB | 0 | 0 | 0 | 0 | 1 | 1 | 0 | 0 | 0 | 0 | 1 | 0 | 3 |
| AvrE | 1 | 1 | 1 | 1 | 1 | 1 | 1 | 2 | 1 | 2 | 1 | 1 | 1 |
| AvrPto | 1 | 0 | 0 | 0 | 1 | 1 | 0 | 0 | 0 | 0 | 0 | 0 | 0 |
| AvrRpm | 0 | 0 | 0 | 0 | 2 | 1 | 0 | 0 | 0 | 0 | 1 | 1 | 0 |
| AvrRpt | 0 | 0 | 1 | 0 | 0 | 0 | 0 | 0 | 0 | 0 | 0 | 0 | 0 |
| HopA | 1 | 1 | 2 | 1 | 0 | 2 | 1 | 0 | 0 | 0 | 0 | 0 | 0 |
| HopB | 3 | 2 | 2 | 1 | 1 | 2 | 1 | 2 | 2 | 1 | 1 | 2 | 2 |
| HopC | 1 | 0 | 2 | 1 | 0 | 0 | 0 | 0 | 0 | 0 | 0 | 0 | 0 |
| HopD | 3 | 1 | 1 | 0 | 2 | 2 | 0 | 0 | 0 | 0 | 0 | 0 | 1 |
| HopE | 1 | 0 | 0 | 0 | 0 | 0 | 0 | 0 | 0 | 0 | 0 | 0 | 0 |
| HopF | 1 | 1 | 1 | 2 | 3 | 3 | 1 | 0 | 0 | 0 | 0 | 1 | 1 |
| HopG | 1 | 0 | 0 | 0 | 0 | 0 | 0 | 0 | 0 | 0 | 0 | 0 | 1 |
| HopH | 1 | 0 | 1 | 0 | 0 | 1 | 0 | 1 | 1 | 1 | 1 | 1 | 0 |
| HopI | 1 | 1 | 1 | 0 | 1 | 1 | 1 | 1 | 1 | 1 | 1 | 1 | 1 |
| HopK | 1 | 1 | 0 | 0 | 0 | 0 | 0 | 0 | 0 | 0 | 0 | 0 | 1 |
| HopL | 1 | 1 | 1 | 1 | 0 | 0 | 1 | 0 | 0 | 0 | 1 | 0 | 0 |
| HopM | 1 | 1 | 2 | 1 | 1 | 1 | 1 | 1 | 1 | 1 | 1 | 1 | 1 |
| HopN1 | 1 | 1 | 0 | 0 | 1 | 1 | 0 | 0 | 0 | 0 | 0 | 0 | 0 |
| HopO | 5 | 3 | 5 | 1 | 0 | 0 | 2 | 0 | 0 | 0 | 0 | 0 | 0 |
| HopQ | 1 | 1 | 1 | 0 | 1 | 1 | 0 | 0 | 0 | 0 | 0 | 0 | 1 |
| HopR | 1 | 1 | 1 | 1 | 1 | 1 | 1 | 0 | 0 | 0 | 0 | 0 | 1 |
| HopS | 1 | 1 | 1 | 2 | 1 | 1 | 1 | 0 | 0 | 0 | 0 | 0 | 0 |
| HopT | 2 | 1 | 2 | 0 | 0 | 0 | 1 | 0 | 0 | 0 | 0 | 0 | 0 |
| HopU | 1 | 1 | 0 | 0 | 0 | 0 | 0 | 0 | 0 | 0 | 0 | 0 | 0 |
| HopV | 1 | 0 | 0 | 2 | 0 | 1 | 0 | 0 | 0 | 0 | 0 | 0 | 1 |
| HopW | 0 | 0 | 2 | 1 | 3 | 2 | 0 | 1 | 1 | 1 | 1 | 1 | 3 |
| HopX | 1 | 1 | 0 | 0 | 1 | 0 | 1 | 0 | 0 | 0 | 1 | 0 | 1 |
| HopY | 1 | 1 | 1 | 3 | 1 | 1 | 1 | 0 | 0 | 0 | 0 | 0 | 0 |
| HopZ | 0 | 0 | 0 | 0 | 0 | 1 | 0 | 0 | 0 | 0 | 0 | 1 | 0 |
| HopAA | 2 | 1 | 1 | 1 | 1 | 3 | 1 | 4 | 1 | 4 | 1 | 1 | 2 |
| HopAB | 1 | 1 | 1 | 0 | 1 | 0 | 1 | 2 | 1 | 2 | 1 | 0 | 3 |
| HopAD | 1 | 0 | 0 | 0 | 0 | 0 | 0 | 0 | 0 | 0 | 0 | 0 | 0 |
| HopAF | 1 | 1 | 1 | 3 | 1 | 1 | 1 | 0 | 0 | 0 | 1 | 1 | 1 |
| HopAG | 2 | 1 | 1 | 2 | 1 | 2 | 1 | 1 | 1 | 1 | 1 | 2 | 0 |
| HopAH | 3 | 3 | 3 | 0 | 3 | 3 | 3 | 1 | 2 | 1 | 2 | 2 | 1 |
| HopAI | 1 | 1 | 1 | 2 | 2 | 1 | 1 | 2 | 1 | 2 | 1 | 2 | 0 |
| HopAL | 0 | 0 | 0 | 0 | 0 | 0 | 0 | 1 | 1 | 1 | 0 | 1 | 0 |
| HopAM | 2 | 0 | 0 | 0 | 1 | 1 | 0 | 0 | 0 | 0 | 0 | 0 | 0 |
| HopAQ | 1 | 0 | 0 | 0 | 0 | 0 | 0 | 0 | 0 | 0 | 0 | 0 | 0 |
| HopAR | 0 | 0 | 0 | 0 | 1 | 0 | 0 | 0 | 1 | 1 | 0 | 0 | 0 |
| HopAS | 2 | 2 | 1 | 2 | 1 | 1 | 1 | 0 | 0 | 0 | 0 | 0 | 1 |
| HopAT | 2 | 0 | 1 | 0 | 1 | 2 | 0 | 0 | 0 | 0 | 0 | 0 | 3 |
| HopAU | 0 | 0 | 0 | 0 | 1 | 1 | 0 | 0 | 0 | 0 | 0 | 0 | 1 |
| HopAW | 0 | 0 | 0 | 0 | 0 | 1 | 0 | 0 | 0 | 0 | 0 | 0 | 1 |
| HopAZ | 0 | 0 | 0 | 1 | 1 | 1 | 0 | 0 | 0 | 0 | 0 | 0 | 0 |
| HopBC | 0 | 0 | 0 | 0 | 0 | 0 | 0 | 0 | 0 | 0 | 0 | 1 | 0 |
| HopBE | 0 | 0 | 0 | 0 | 0 | 0 | 0 | 0 | 0 | 0 | 0 | 1 | 0 |
| HopBK | 0 | 0 | 0 | 0 | 0 | 0 | 0 | 0 | 0 | 1 | 0 | 0 | 0 |
| HopBM | 2 | 0 | 0 | 0 | 0 | 0 | 0 | 0 | 0 | 0 | 0 | 0 | 0 |
| HopBN | 0 | 0 | 1 | 2 | 1 | 1 | 0 | 0 | 0 | 0 | 0 | 0 | 0 |
| HopBO | 0 | 0 | 0 | 0 | 1 | 0 | 1 | 0 | 0 | 0 | 0 | 0 | 0 |
| HopBP | 0 | 0 | 0 | 0 | 1 | 1 | 2 | 0 | 0 | 0 | 1 | 0 | 0 |
| HopBQ | 0 | 0 | 0 | 0 | 2 | 0 | 0 | 0 | 0 | 0 | 0 | 0 | 0 |

**Supplementary Table S3. Known interaction outcomes of *P. syringae* WT and *avrB*-expressing strains in *Arabidopsis* *thaliana* Col-0.**

| **Phylogroup** | **Species** | **Strain** | **Interaction outcome**  **WT strain / *A. thaliana* Col-0** | **Interaction outcome**  ***avrB-*expressing strain / *A. thaliana* Col-0** |
| --- | --- | --- | --- | --- |
| 1a | *P*. *syringae* pv. *tomato* | DC3000 | Compatible/disease (no HR)  (Preston, 2000; Xin and He, 2013) | Avirulent/host resistance (HR)  (Innes et al., 1993; Mackey et al., 2002; Mudgett, 2005) |
|  | *P*. *syringae* pv. *maculicola* | M6 | Compatible/disease (no HR)  (Debener et al., 1991) | Avirulent/host resistance (HR)  (This study) |
|  | *P*. *syringae* pv. *tomato* | T1 | Non-host resistance type II (HR)  (Ishiga et al., 2011) | Non-host resistance type II (HR)  (This study) |
|  | *P*. *syringae* pv. *actinidiae* | CRA-FRU 8.43 | Non-host resistance type I (no HR)  (Jayaraman et al., 2017) | Non-host resistance type II (HR)  (Puttilli et al., 2022) |
| 2d | *P*. *syringae* pv. *aptata* | CC0094 | Non-host resistance type II (HR)  (This study) | Non-host resistance type II (HR)  (This study) |
|  | *P*. *syringae* pv. *syringae* | B728a | Non-host resistance type I (no HR)  (Vinatzer et al., 2006) | Non-host resistance type II (HR)  (This study) |
| 3a | *P*. *syringae* pv. *phaseolicola* | 1448A | Non-host resistance type I (no HR)  (Ham et al., 2007) | Non-host resistance type II (HR)  (This study) |
